# ACE2 and TMPRSS2 are expressed on the human ocular surface, suggesting susceptibility to SARS-CoV-2 infection

**DOI:** 10.1101/2020.05.09.086165

**Authors:** Lingli Zhou, Zhenhua Xu, Gianni M. Castiglione, Uri S. Soiberman, Charles G. Eberhart, Elia J. Duh

## Abstract

**Purpose:** Conjunctival signs and symptoms are observed in a subset of patients with COVID-19, and SARS-CoV-2 has been detected in tears, raising concerns regarding the eye both as a portal of entry and carrier of the virus. The purpose of this study was to determine whether ocular surface cells possess the key factors required for cellular susceptibility to SARS-CoV-2 entry/infection.

**Methods:** We analyzed human post-mortem eyes as well as surgical specimens for the expression of ACE2 (the receptor for SARS-CoV-2) and TMPRSS2, a cell surface-associated protease that facilitates viral entry following binding of the viral spike protein to ACE2.

**Results:** Across all eye specimens, immunohistochemical analysis revealed expression of ACE2 in the conjunctiva, limbus, and cornea, with especially prominent staining in the superficial conjunctival and corneal epithelial surface. Surgical conjunctival specimens also showed expression of ACE2 in the conjunctival epithelium, especially prominent in the superficial epithelium, as well as the substantia propria. All eye and conjunctival specimens also expressed TMPRSS2. Finally, western blot analysis of protein lysates from human corneal epithelium obtained during refractive surgery confirmed expression of ACE2 and TMPRSS2.

**Conclusions:** Together, these results indicate that ocular surface cells including conjunctiva are susceptible to infection by SARS-CoV-2, and could therefore serve as a portal of entry as well as a reservoir for person-to-person transmission of this virus. This highlights the importance of safety practices including face masks and ocular contact precautions in preventing the spread of COVID-19 disease.

## 1. Introduction

With the COVID-19 pandemic, there has been heightened awareness and interest in the routes of viral infection, given both the public health and medical implications. Conjunctival abnormalities are a recently documented manifestation of COVID-19 [1]. In addition, various investigators have reported the presence of virus in tear and conjunctival swab specimens in a subset of patients with COVID-19 [2, 3]. Consequently, there has been extensive speculation regarding the ocular surface as a possible site of virus entry and also as a source of contagious infection [4, 5]. The ocular surface could potentially serve as a portal of entry through exposure to aerosolized droplets or hand-eye contact. Similarly, the ocular surface could serve as an important reservoir of virus that could result in transmission to other individuals. In addition to issues related to transmission, ocular surface susceptibility to infection has implications for ophthalmic manifestations of COVID-19.

Recent studies of the respiratory tract have confirmed the critical elements for cellular susceptibility to infection by SARS-CoV-2. Notably, nasal epithelial cells and bronchial secretory cells are known to possess the key factors for cellular susceptibility to SARS-CoV-2 [6, 7]. ACE2 serves as the key cell-surface receptor for SARS-CoV-2 [8-12] that binds the viral spike protein [10-14], and TMPRSS2 is known to be an important cell surface-associated protease that allows viral entry following binding of the viral spike protein to ACE2 [9, 15]. However, whether ocular surface cells possess these key factors for cellular susceptibility to viral infection remains unclear.

In the current study, we analyzed both human post-mortem whole eyes and surgical conjunctiva specimens using immunohistochemistry in order to determine both expression and localization of these key viral susceptibility factors on the ocular surface. Across all specimens, we found expression of ACE2 in the conjunctiva, limbus, and cornea. All eye and conjunctival specimens also expressed TMPRSS2.

Immunoblotting using the same antibodies further confirmed expression of ACE2 on the human ocular surface. Together, these results indicate that ocular surface cells including conjunctiva and cornea are indeed susceptible to infection by SARS-CoV-2, highlighting the importance of safety practices protecting this region [4].

## 2. Methods

### 2.1. Patients and specimens

Corneal epithelial samples were obtained from healthy myopic patients undergoing photorefractive keratectomy for the treatment of mild to moderate myopia or myopic astigmatism. Topical proparacaine and 5% betadine solution were applied to the ocular surface. The central 9 mm of the corneal epithelium was removed with sterile cellulose sponges and immediately placed in RNAlater® tubes (Thermo Fisher, MA, USA). The samples were kept at -80°C until protein extraction using the PARIS™ kit (Thermo Fisher, Waltham, MA, USA). Protein concentration was quantified by BCA assay (Thermo Fisher). Ten post-mortem human globes were collected from five non-diabetic controls lacking ocular disease, and from five diabetic individuals with diabetic retinopathy. Five surgical conjunctival specimens were obtained during routine eye surgery as described [16]. The tissues were fixed in 10% formalin and embedded in paraffin. Clinical details pertaining to study patients are described in Table 1. Research followed the tenets of the Declaration of Helsinki. Study protocols were approved by the Institutional Review Board at the Johns Hopkins School of Medicine, and study participants underwent informed consent by the treating surgeon.

**Table 1.**
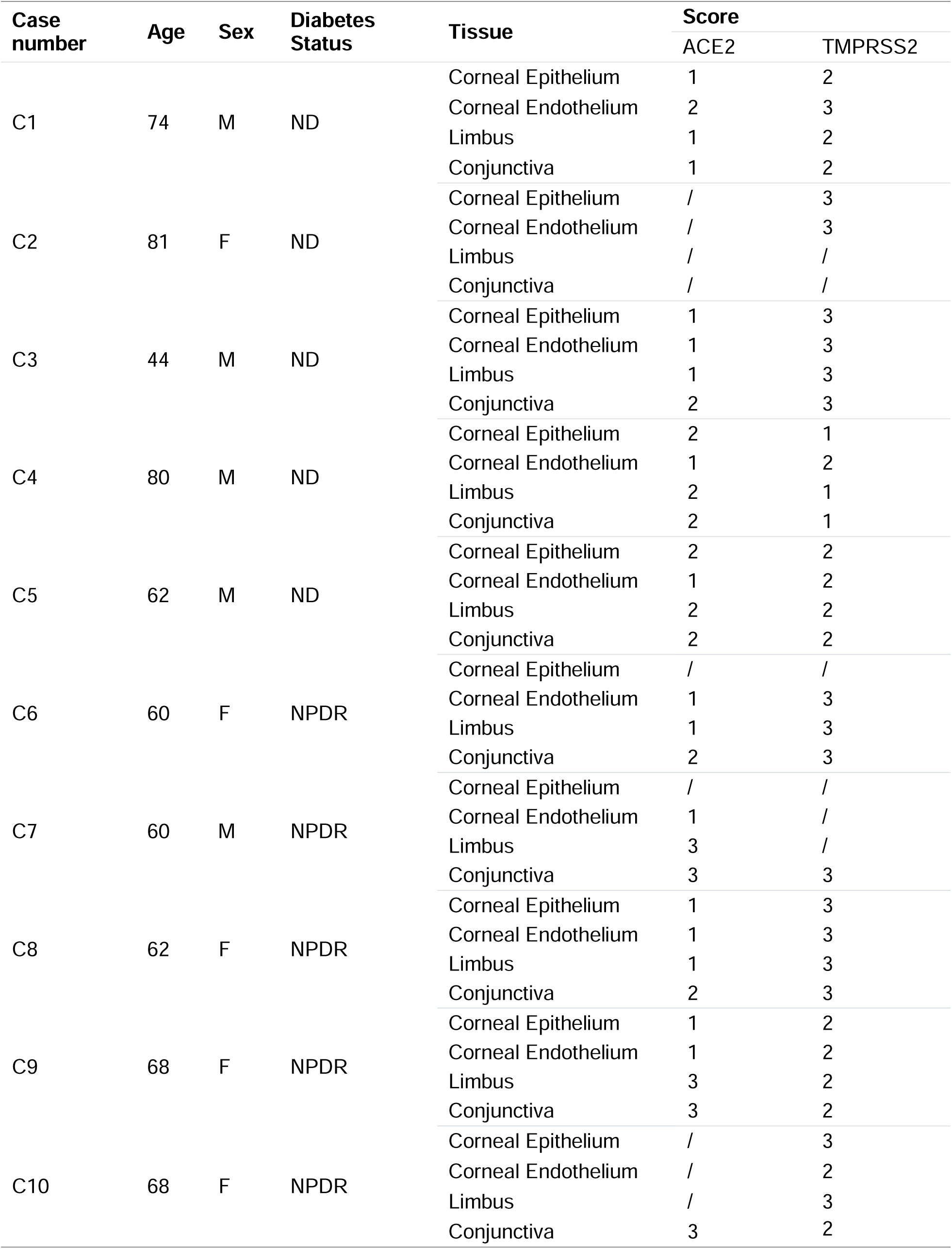
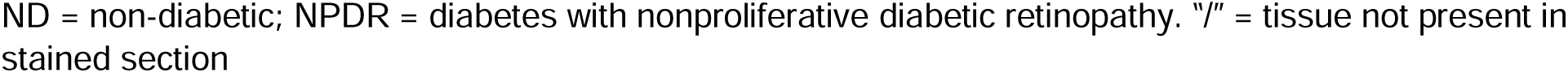
Demographic information and grading score for postmortem eyes.

| Case number | Age | Sex | Diabetes Status | Tissue | Score |  |
| --- | --- | --- | --- | --- | --- | --- |
|  |  |  |  |  | ACE2 | TMPRSS2 |
| C1 | 74 | M | ND | Corneal Epithelium | 1 | 2 |
|  |  |  |  | Corneal Endothelium | 2 | 3 |
|  |  |  |  | Limbus | 1 | 2 |
|  |  |  |  | Conjunctiva | 1 | 2 |
| C2 | 81 | F | ND | Corneal Epithelium | / | 3 |
|  |  |  |  | Corneal Endothelium | / | 3 |
|  |  |  |  | Limbus | / | / |
|  |  |  |  | Conjunctiva | / | / |
| C3 | 44 | M | ND | Corneal Epithelium | 1 | 3 |
|  |  |  |  | Corneal Endothelium | 1 | 3 |
|  |  |  |  | Limbus | 1 | 3 |
|  |  |  |  | Conjunctiva | 2 | 3 |
| C4 | 80 | M | ND | Corneal Epithelium | 2 | 1 |
|  |  |  |  | Corneal Endothelium | 1 | 2 |
|  |  |  |  | Limbus | 2 | 1 |
|  |  |  |  | Conjunctiva | 2 | 1 |
| C5 | 62 | M | ND | Corneal Epithelium | 2 | 2 |
|  |  |  |  | Corneal Endothelium | 1 | 2 |
|  |  |  |  | Limbus | 2 | 2 |
|  |  |  |  | Conjunctiva | 2 | 2 |
| C6 | 60 | F | NPDR | Corneal Epithelium | / | / |
|  |  |  |  | Corneal Endothelium | 1 | 3 |
|  |  |  |  | Limbus | 1 | 3 |
|  |  |  |  | Conjunctiva | 2 | 3 |
| C7 | 60 | M | NPDR | Corneal Epithelium | / | / |
|  |  |  |  | Corneal Endothelium | 1 | / |
|  |  |  |  | Limbus | 3 | / |
|  |  |  |  | Conjunctiva | 3 | 3 |
| C8 | 62 | F | NPDR | Corneal Epithelium | 1 | 3 |
|  |  |  |  | Corneal Endothelium | 1 | 3 |
|  |  |  |  | Limbus | 1 | 3 |
|  |  |  |  | Conjunctiva | 2 | 3 |
| C9 | 68 | F | NPDR | Corneal Epithelium | 1 | 2 |
|  |  |  |  | Corneal Endothelium | 1 | 2 |
|  |  |  |  | Limbus | 3 | 2 |
|  |  |  |  | Conjunctiva | 3 | 2 |
| C10 | 68 | F | NPDR | Corneal Epithelium | / | 3 |
|  |  |  |  | Corneal Endothelium | / | 2 |
|  |  |  |  | Limbus | / | 3 |
|  |  |  |  | Conjunctiva | 3 | 2 |

ND = non-diabetic; NPDR = diabetes with nonproliferative diabetic retinopathy. “/” = tissue not present in stained section

### 2.2. Western blot analysis

Proteins from corneal epithelial samples were extracted using the PARIS kit (Thermo Fisher). Total protein concentration was measured using DC protein assay kit (500-0112, Bio-Rad). 20 µg protein from cell lysates was subjected to 7.5 % SDS-PAGE and transferred to Hybond ECL nitrocellulose membrane (Amersham Biosciences, Piscataway, NJ). After incubation with appropriate primary and secondary antibodies, the blots were detected with the Supersignal Femto Chemiluminescent Substrates (ThermoFisher, MA, USA). For re-probing, blots were washed in Western blot stripping buffer (Thermo Fisher) for 15 minutes before proceeding with new blotting. Anti-ACE2 (ab108252, 1:1000) and anti-TMPRSS2 (ab92323, 1:1000) antibodies were purchased from Abcam. β-actin detected by its antibody (#4970, 1:2000, Cell Signaling) was used for loading control. Horseradish peroxidase-tagged secondary anti-rabbit IgG was from Cell Signaling (#7074, 1:2000).

### 3.3. Immunohistochemistry

Deparaffinized sections were boiled in 1× target retrieval solution (Dako, Capenteria, CA) for 20 min. After blocking in 5% normal goat serum diluted in PBS, the sections were incubated with anti-ACE2 antibody (ab108252, 1: 100, Abcam, Cambridge, MA), anti-TMPRSS2 antibody (ab92323, 1: 1000, Abcam) and the isotype- and concentration-matched normal IgG control (ab172730, Abcam) overnight at 4 ° C. After 3 washes with PBST (PBS plus 0.1% Tween-20), the sections were incubated with the secondary biotinylated goat anti-rabbit IgG (1:2000) for 1 hour at room temperature. The sections were then detected by alkaline phosphatase detection system (Vectastain ABC-AP kit, Vector laboratories, Burlingame, CA) and a blue reaction product was produced by incubating sections with alkaline substrate (Vector blue AP substrate kit III, Vector laboratories) for 20 to 40 minutes. The sections were also counterstained with nuclear fast red (Vector laboratories) and mounted with mounting medium (H-5501, Vector laboratories), and examined under a light microscope.

### Grading of immunohistochemistry

ACE2 and TMPRSS2 expression levels in the ocular surface tissues were scored based on the relative intensity of ACE2 and TMPRSS2 immunoreactivity. The intensity was graded by two independent observers in a masked fashion. The grading scores for intensity were: 0, no staining (similar to non-immune IgG -incubated control staining); 1, mild staining; 2, moderate staining and 3, strong staining. For this grading, immunohistochemistry was performed at the same time for all post-mortem globe, or surgical conjunctiva sections. For some post-mortem globe specimens, conjunctiva, limbus, and/or cornea were not present in the stained sections, as indicated in Table 1.

## 3. Results

### 3.1. ACE2 and TMPRSS2 are expressed in the human corneal epithelium

Several studies have demonstrated ACE2 expression in epithelial cells of tissues including lung, ileum and kidney. In order to determine whether ACE2 is also expressed in corneal epithelial cells, we performed western blot analysis of corneal epithelium obtained from two patients during photorefractive keratectomy. Western blot analysis clearly showed that ACE2 protein was expressed in both corneal epithelial lysates as a single 110 kDa band (Fig. 1), which represents the glycosylated form of ACE2 [17]. TMPRSS2 is known to be a key protein for priming the SARS-CoV-2 Spike protein for host membrane fusion to enable cell entry. Western blot data similarly showed the expression of TMPRSS2 protein in the human corneal epithelial samples (Fig. 1). Taken together, ACE2 and TMPRSS2 protein were both expressed in corneal epithelial cells.

**Fig. 1.**
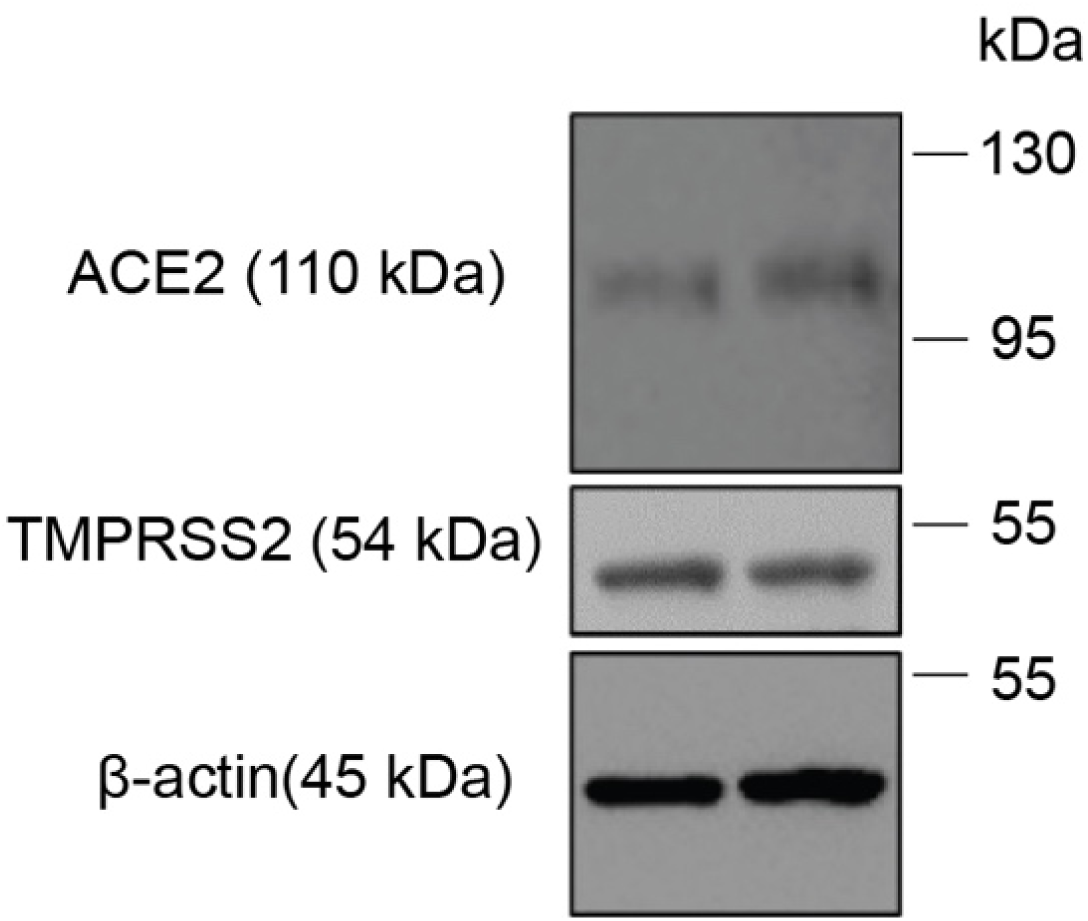
ACE2 and TMPRSS2 protein expression in human corneal epithelium. Tissue lysates were prepared from corneal epithelial cell samples collected from two patients and analyzed by Western blot, using antibodies against ACE2 or TMPRSS2. Beta-actin was used as the loading control.

### 3.2. Expression and localization of ACE2 in cornea and limbus

Using the same antibody, we further confirmed the expression of ACE2 and investigated its localization in cornea by immunohistochemical staining of ten post-mortem eye specimens (Table 1). Consistent with our western blot analysis, ACE2 staining was positive in corneal epithelium, with mild to moderate staining in both basal cells and flattened ones in the most superficial layer (Fig. 2A and B). ACE2 staining was largely absent in the corneal stroma (Fig. 2A). There was ACE2 expression in the corneal endothelium as well, with staining intensity roughly equivalent to corneal epithelium (Fig. 2C). The corneal limbus was also positive for ACE2 staining (Fig. 2E), with particularly prominent staining in the superficial epithelial cells (Fig. 2F). In some cases, ACE2 staining intensity in the limbus was stronger than the adjacent corneal epithelium.

**Fig. 2.**
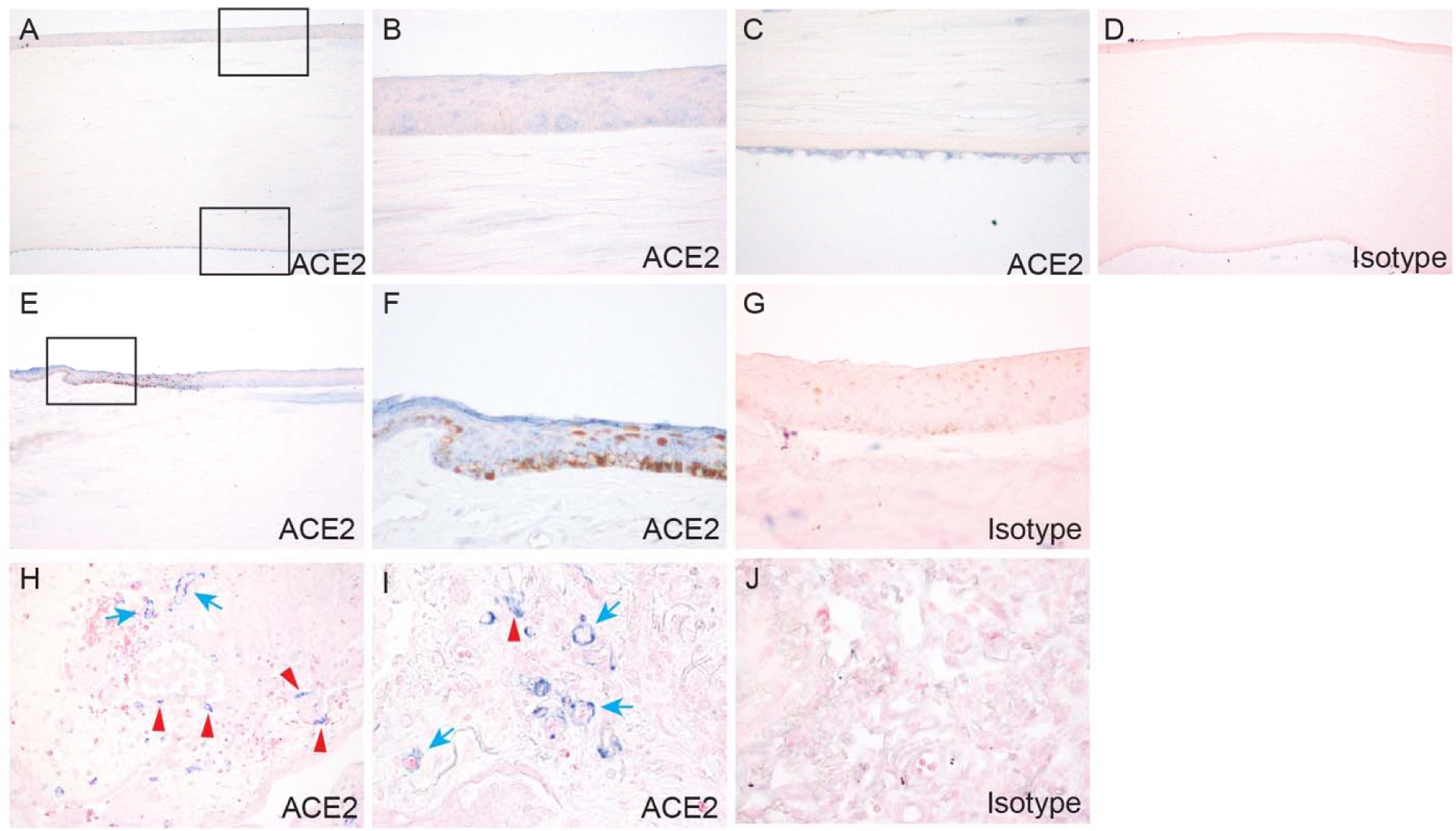
Expression and localization of ACE2 in cornea and limbus. Post-mortem human eye sections were processed for immunohistochemistry with rabbit anti-ACE2 antibody. ACE2 expression was analyzed in cornea (A-C), limbus (E-F), and lung (H-I). Boxed region in (A) is shown at higher magnification in (B, C). Boxed region in (E) is shown at higher magnification in (F). Lung epithelium and vascular endothelium are indicated by red and blue arrow respectively. Immunohistochemistry with isotype- and concentration-matched normal IgG control was performed (D, G, J).

As an additional positive control, we analyzed lung tissue with the same ACE2 antibody. Consistent with published reports, ACE2 was expressed in both a subset of lung epithelial cells, and many vascular endothelial cells (Fig. 2H and I) [18, 19]. For both corneal and lung sections, immunohistochemical staining with isotype IgG isotype antibody staining was performed, as negative controls (Fig. 2D, G and J).

### 3.3. Expression and localization of ACE2 in conjunctiva

The conjunctiva occupies the majority of the ocular surface and has been speculated to represent a significant target for SARS-CoV-2 infection. As shown in Fig. 3, ACE2 was indeed expressed in conjunctival epithelium. In many specimens, ACE2 staining was especially prominent in the most superficial epithelial cells (Fig. 3A, B), while in others a mixture of superficial and basal epithelial cells were stained (Fig. 3C, D). Goblet cells were largely negative (Fig. 3D, arrows). In terms of subcellular localization, ACE2 was mainly detected in cell membranes and the cytoplasm, with no nuclear staining. Beneath the epithelium, there was very little or no ACE2 staining in the substantia propria, and unlike the lung vascular endothelial cells were negative (Fig. 3A, arrow). Among all globe specimens examined, moderate to strong ACE2 expression was most frequently identified (Table 1), and the distribution of staining was similar.

**Fig. 3.**
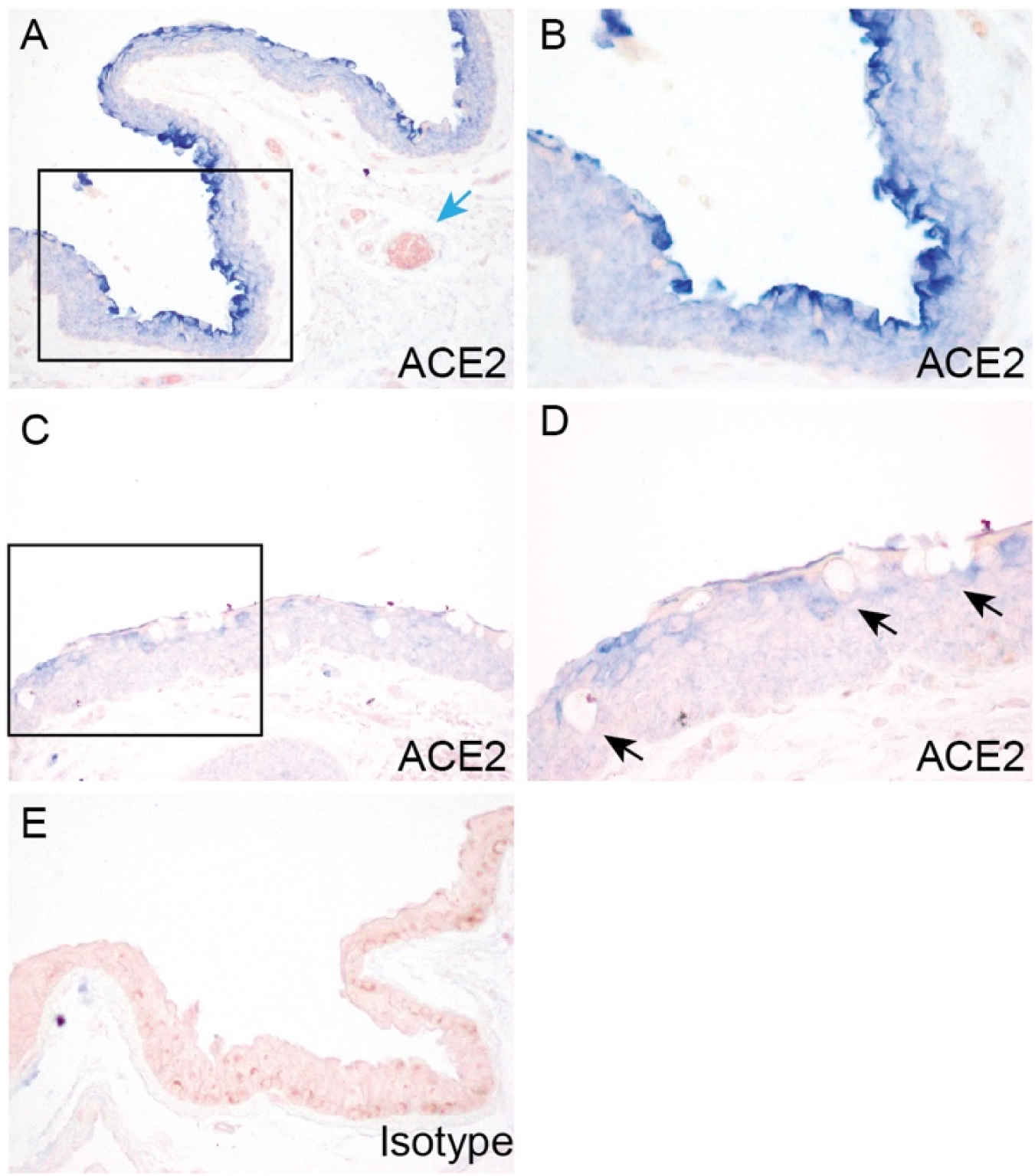
Expression and localization of ACE2 in conjunctiva in post-mortem globe. Paraffin-embedded human eye sections were processed for immunohistochemistry with rabbit anti-ACE2 antibody. ACE2 expression was analyzed in conjunctiva. Conjunctiva from two separate globes are depicted (A-B, C-D). For each individual globe, boxed region is shown at higher magnification. Arrows indicate vascular endothelium (A) and goblet cells (D). Immunohistochemistry with isotype- and concentration-matched normal IgG control was performed (E).

As a complementary study to the post-mortem globe analysis, we also performed immunohistochemical analysis of surgical conjunctival specimens (Table 2). ACE2 expression was positive across all five specimens, and the expression pattern was similar to what we observed in post-mortem eye specimens (Fig. 4A-E). ACE2 was most strongly expressed in apical conjunctival epithelium, with variable expression in more basal cells, and weak or focal expression in the substantia propria (Fig. 4A-E). Goblet cells in epithelium were negative for ACE2 staining (Fig. 4C), as were vascular endothelium. We also found ACE2 staining was positive in conjunctival invaginations and pseudoglands of Henle (Fig. 4D and E). Immunohistochemical staining with isotype IgG isotype antibody staining was performed, as negative control (Fig. 4F).

**Table 2.** Demographic information and grading score for surgical conjunctival specimens.

| Case number | Age | Sex | Score |  |
| --- | --- | --- | --- | --- |
|  |  |  | ACE2 | TMPRSS2 |
| SC 1 | 84 | M | 3 | 3 |
| SC 2 | 41 | M | 2 | 3 |
| SC 3 | 70 | F | 2 | 3 |
| SC 4 | 41 | M | 1 | 2 |
| SC 5 | 53 | F | 3 | 2 |

**Fig. 4.**
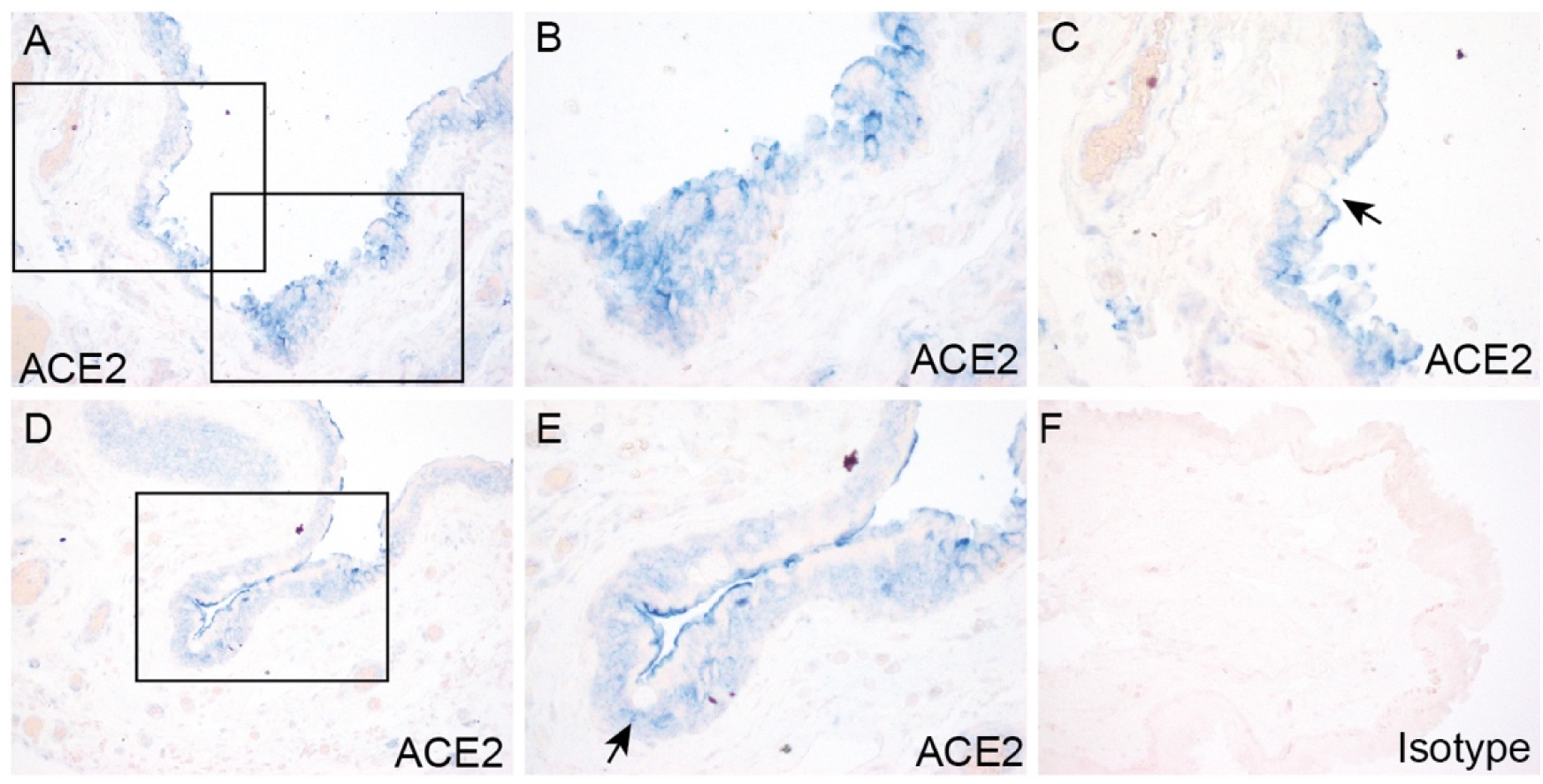
Expression and localization of ACE2 in surgical conjunctival specimens. Paraffin-embedded human surgical conjunctival sections were analyzed by immunohistochemistry with rabbit anti-ACE2 antibody. Boxed regions in (A) are shown at higher magnification in (B) and (C). Boxed region in (D) is shown at higher magnification in (E). Arrows indicate goblet cells (C, E). Immunohistochemistry with isotype- and concentration-matched normal IgG control was performed (F).

### 3.4. Expression and localization of TMPRSS2 in cornea and limbus

We next examined the expression and localization of TMPRSS2 in corneas of the post-mortem eyes by immunohistochemical staining. In cornea, TMPRSS2 uniformly exhibited very strong staining in both epithelium and endothelium (Fig. 5A-C). In contrast to ACE2, in corneal epithelium TMPRSS2 was expressed in all layers, including superficial flattened cells, middle wing cells, and deep basal cells. TMPRSS2 staining was also weakly positive in some keratocytes of corneal stroma. In the limbus, TMPRSS2 exhibited intense staining in the multi-layered epithelium (Fig. 5E and F). As a positive control, we analyzed lung tissue with the same TMPRSS2 antibody. TMPRSS2 showed positive staining in a broader range of lung cells as compared to ACE2 (Fig. 5H). For both corneal and lung sections, immunohistochemical staining with isotype IgG isotype antibody staining was performed, as negative controls (Fig. 5D, G and J).

**Fig. 5.**
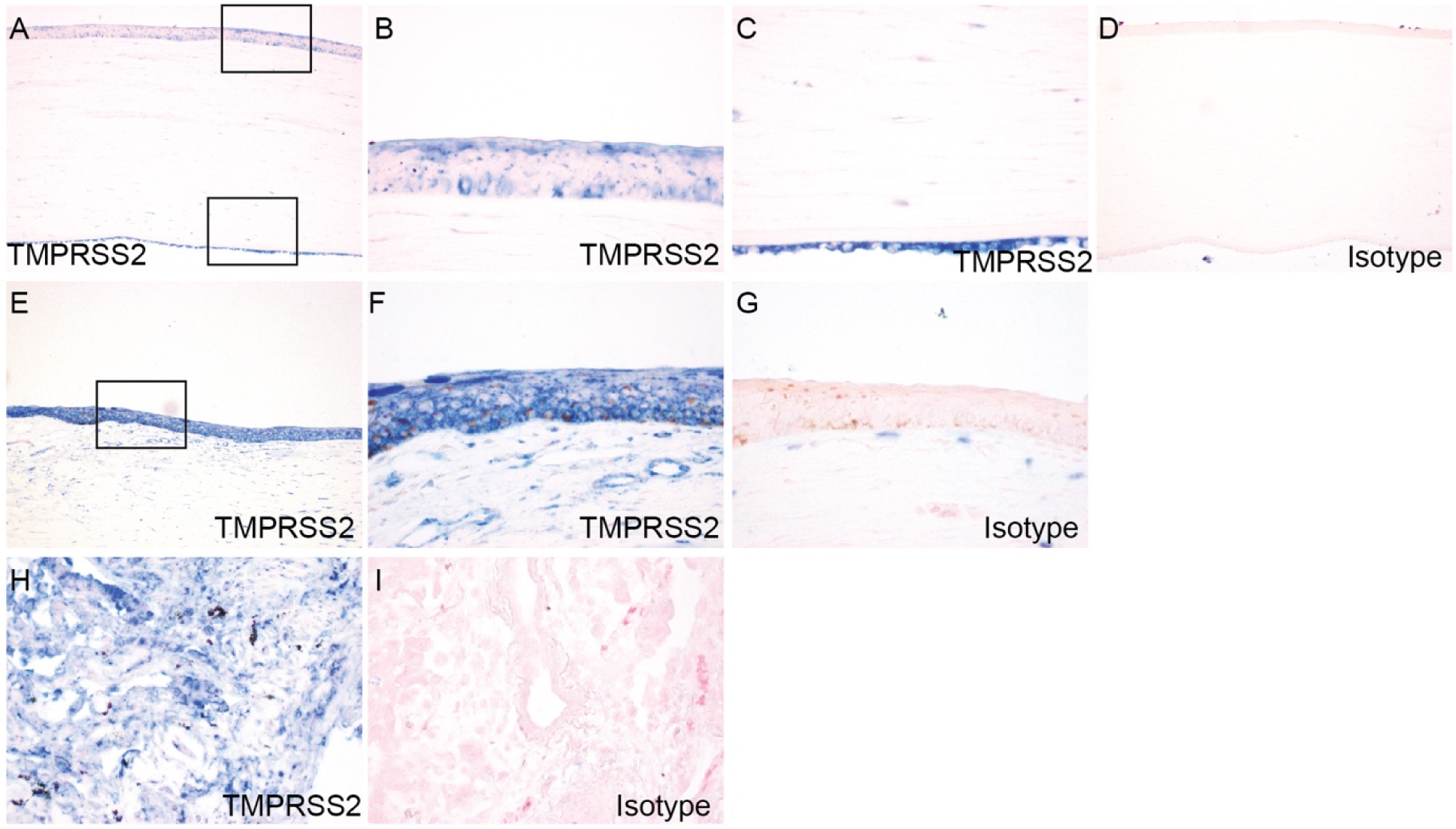
Expression and localization of TMPRSS2 in cornea and limbus. Paraffin-embedded human eye sections were processed for immunohistochemistry with rabbit anti-TMPRSS2 antibody. ACE2 expression was analyzed in cornea (A-C), limbus (E-F), and lung (H). Boxed region in (A) is shown at higher magnification in (B, C). Boxed region in (E) is shown at higher magnification in (F). Immunohistochemistry with isotype- and concentration-matched normal IgG control was performed (D, G, I).

### 3.5. Expression and localization of TMPRSS2 in conjunctiva

In conjunctiva from post-mortem eye specimens, TMPRSS2 was expressed throughout the whole epithelial layer (Fig. 6A-F), in contrast to ACE2 staining which was more prominent in the apical epithelium. In some specimens, TMPRSS2 showed stronger staining in basal cells compared with other layers of conjunctival epithelium (Fig. 6E and F). Similar to ACE2, TMPRSS2 staining was negative in goblet cells (Fig. 6D). In the corneal stroma, TMPRSS2 showed mild staining in fibroblasts and vascular endothelium (Fig. 6A).

**Fig. 6.**
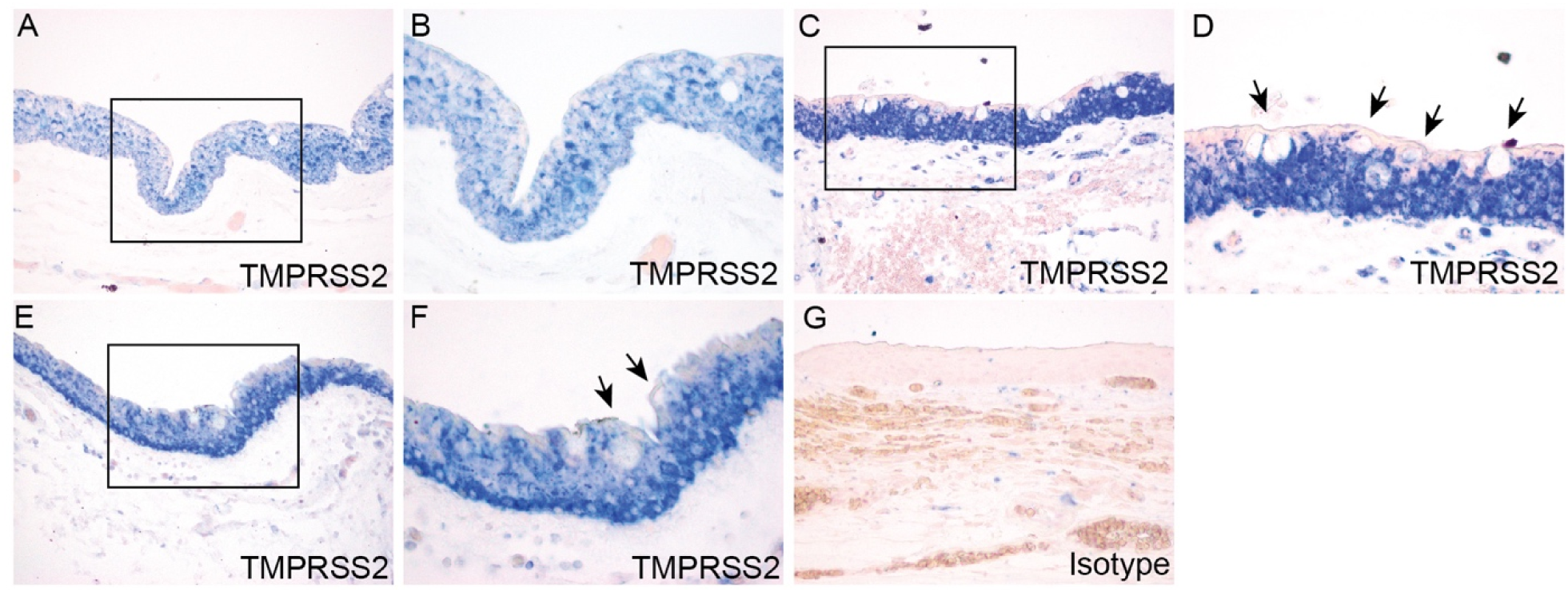
Expression and localization of TMPRSS2 in conjunctiva in post-mortem globe. Paraffin-embedded human eye sections were processed for immunohistochemistry with rabbit anti-TMPRSS2 antibody. ACE2 expression was analyzed in conjunctiva. Conjunctiva from three separate globes are depicted (A-B, C-D, E-F). For each individual globe, boxed region is shown at higher magnification. Arrows indicate goblet cells (D, F). Immunohistochemistry with isotype- and concentration-matched normal IgG control was performed (G).

As a complementary study to the post-mortem globe analysis, we also performed immunohistochemical studies of surgical conjunctival specimens (Table 2). In all five surgical conjunctiva specimens, TMPRSS2 staining was positive throughout the epithelium (Fig. 7), with a similar staining pattern as compared with conjunctiva from post-mortem eye specimens.

**Fig. 7.**
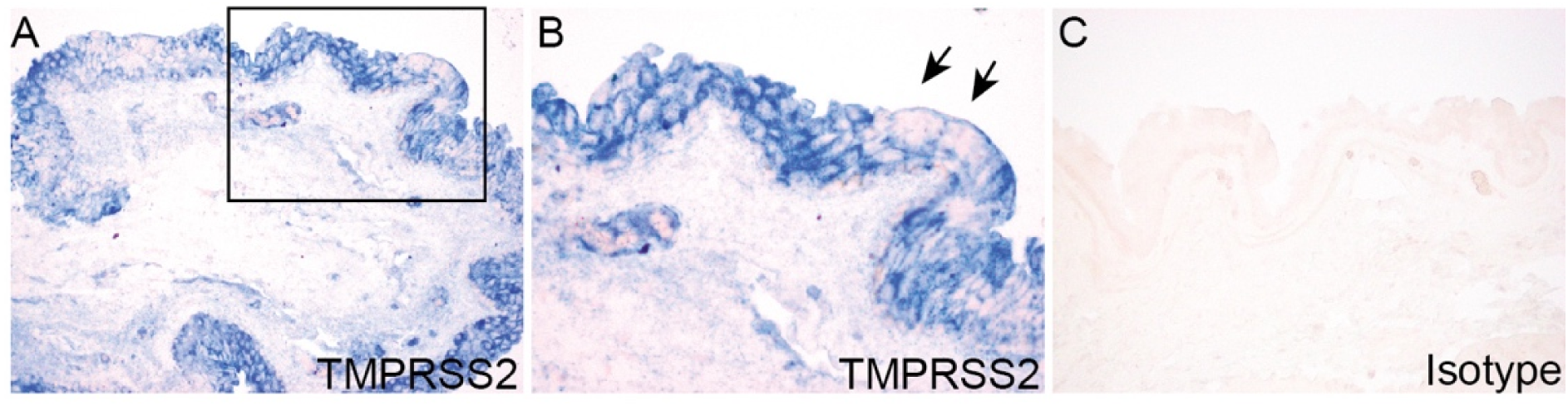
Expression and localization of TMPRSS2 in surgical conjunctival specimens. Paraffin-embedded human surgical conjunctival sections were analyzed by immunohistochemistry with rabbit anti-TMPRSS2 antibody. Boxed region in (A) is shown at higher magnification in (B). Arrows indicate goblet cells (B). Immunohistochemistry with isotype- and concentration-matched normal IgG control was performed (C).

### 3.6. Grading of ACE2 and TMPRSS2 expression in epithelium on the ocular surface

As an additional analysis, we graded the staining intensities of ACE2 and TMPRSS2 on the ocular surface from 10 post-mortem eye specimens with particular attention to their expression in epithelium. For some post-mortem globe specimens, conjunctiva, limbus, and/or cornea were not retained for some stained sections. As shown in Table 1, ACE2 was expressed in the epithelium of cornea, limbus, and conjunctiva in all intact specimens to varying degrees. Overall, ACE2 exhibited much stronger staining in conjunctival epithelium than the epithelium from cornea and limbus. TMPRSS2 exhibited consistent strong staining in the epithelial layers of cornea, limbus, and conjunctiva in all specimens. ACE2 and TMPRSS2 both were expressed in the corneal endothelium from all 10 specimens. TMPRSS2 exhibited more intensive staining compared to ACE2. We further graded the staining intensities of ACE2 and TMPRSS2 in surgical conjunctiva specimens (Table 2). In comparison to post-mortem globe sections, surgical conjunctival specimens exhibited greater intensity of ACE2 staining. TMPRSS2 showed strong staining in most specimens, whether globe or surgical conjunctiva.

## 4. Discussion

An important aspect of the ongoing efforts to control and mitigate the impact of the COVID-19 pandemic has been to reduce the transmission of virus between individuals. In this respect, it is vital to gain greater understanding of the routes and modes of transmission, including the role of the ocular surface. Although the expression of the key cellular elements for susceptibility to infection, including the ACE2 receptor and the membrane-associated TMPRS2, have been well-studied in cells of the respiratory tract, there have been minimal reports to date relating to the eye. In our study, we found expression of both ACE2 and TMPRSS2 across all human ocular specimens. Staining of ACE2 was especially prominent in the most superficial epithelium in both the conjunctiva and cornea. We also found TMPRSS2 in conjunctiva and cornea, with a more ubiquitous staining pattern as compared to ACE2. These results indicate that the ocular surface is indeed susceptible to infection by SARS-CoV-2. The prominent expression of ACE2 on the most superficial corneal and conjunctival epithelium, the site of greatest exposure, is particularly notable.

Several clinical studies have reported the presence of SARS-CoV-2 in tear specimens from individuals with COVID-19. There has been discordance in reports regarding the proportion of COVID-19 patients with presence of virus in ocular specimens, possibly relating to factors including sensitivity of tests, type of ocular specimen and timing, type of specimen obtained, and timing of specimen procurement during disease progression in patients [20]. Studies have also indicated the variable presentations of COVID-19 disease. For instance, conjunctivitis is occasionally the initial clinical presentation of COVID-19 [21]. In addition, presence of virus in ocular swabs have indicated presence of virus in the eye as long as 27 days after initial symptoms, even with concomitant absence of virus in nasal swabs [22]. Although viral infection of ocular cells has not yet been reported in patients, a recent report found SARS-CoV-2 can infect conjunctival epithelium in an ex-vivo culture system [23]. Our results may offer additional insights regarding the role of the ocular surface in COVID-19 transmission, suggesting that the ocular surface may serve as an even more significant reservoir for virus than suggested by current clinical studies.

A very recent study also investigated ACE2 expression in human conjunctiva. Strikingly, in contrast to our study, this report did not find evidence of significant conjunctival ACE2 expression [24]. The explanation for the contrast in our findings is unclear, but might include methodological differences in the immunohistochemistry procedure, quality of tissue specimens, and differences in the antibodies used. In our study, we corroborated IHC studies of ACE2 and TMPRSS2 expression in with western blot analysis using the same antibodies as for IHC. Of note, IHC studies using IgG-isotype control demonstrated no staining for the respective proteins.

The susceptibility of the conjunctival to infection with SARS-CoV-2 provides further context for published clinical studies. COVID-19 is often associated with conjunctival manifestations including conjunctivitis, which was present in about 30 percent of one clinical study of COVID-19 patients [1]. Primary viral infection of the conjunctiva could induce a local immune or inflammatory response resulting in these and other ophthalmic manifestations. In addition, our results support viral tropism for corneal epithelial cells and raises the possibility of corneal manifestations. Although conjunctival findings have been predominantly characterized in ocular COVID-19 disease, there have been some reports about corneal abnormalities [21].

In summary, the presence of ACE2 and TMPRSS2 in conjunctival and corneal epithelial cells supports the ocular surface as a secondary site of infection following respiratory tract, or possibly even as the initial portal of entry to an individual. Infection of ocular surface cells could lead to the eye as being an important carrier, with ocular virus shedding constituting a significant mechanism for infection of other individuals. Our study therefore highlights the importance of safety practices in the general community to prevent infection and spread (hygiene, face masks) and need for extra caution among ophthalmologists [25].

## Acknowledgments

This work was supported by research grants from the National Institutes of Health (EY022383 and EY022683; to E.J.D.) and Core Grant P30EY001765, Imaging and Microscopy Core Module. The authors thanks S. Biswal (Johns Hopkins School of Public Health) for post-mortem lung sections.

